# Prior Perceptual-Cognitive Training Builds Mental Resistance During Acute Physical Fatigue In Professional Rugby Athletes

**DOI:** 10.1101/505313

**Authors:** Jocelyn Faubert, Sylvain Barthes

**Affiliations:** Faubert Lab, School of Optometry, University of Montreal

**Keywords:** Visual Perception, Visual Cognition, Dynamic Attention, Decision Making, Sport Science, Fitness, Fatigue, Rugby, 3D-MOT, NeuroTracker

## Abstract

Elite athletes are engaged in intensive physical training programs because it improves their physical strength and agility, but also to build endurance for resisting the negative impact of fatigue. Although it is well accepted that physical fitness can reduce the consequence of mental fatigue it is still unknown whether specific mental training can reduce the negative impact of physical fatigue on perceptual-cognitive functions. Here we show in professional rugby players that acute physical fatigue can dramatically reduce mental performance required for processing 3D-Multiple Object Tracking (3D-MOT) and that prior training, void of any physical demands, can build resistance to these detrimental effects.

## Introduction

It was recently shown that the level of athletic performance is directly related to the capacity for learning complex and neutral dynamic visual scenes void of motor control requirements and sports specific context (3). In that study, athletes from top professional soccer, rugby and ice hockey leagues were compared to high-level amateurs and university students and demonstrated dramatic differences in their capacity for learning 3D-Multiple Object Tracking (3D-MOT) a demanding dynamic attention task (NeuroTracker^™^) (4) as compared to elite amateurs, who were superior to university students (3). This argues that the mental capacity for processing and learning this task is highly relevant to sport performance level. Furthermore, a recent study from a different laboratory using the same 3D-MOT methodology (NeuroTracker^™^) has shown that a preseason measure of the performance on this task was highly predictive of basketball decision-making performance metrics measured throughout the season for a professional NBA team (6). Another recent study has demonstrated that when soccer athletes from a university varsity team were trained on 3D-MOT they showed a 15% increase in passing decision success in short-sided games (evaluated blind) while active and passive control condition groups showed no improvement (7). Given the pertinence of this perceptual-cognitive ability in high-level sports, it is critical to know how physical fatigue influences this mental ability and whether it is possible to palliate this with training.

The impact of physical fatigue on mental performance is generally well accepted in sports, as mental execution errors are often associated with physical fatigue represented at the end of games. Aerobic training has been shown to improve resistance to mental fatigue (8). However, there is no direct evidence that prior mental training can reduce the negative impact of physical fatigue on mental abilities associated with making appropriate play making decisions on the field as has been demonstrated for 3D-MOT. A recent meta-analytical study looking at the impact of acute exercise on speed and accuracy of cognition has shown that acute exercise improves speed of processing due to an assumed increased arousal (2) but the impact on accuracy measures was inconclusive (9). They state that, “The very limited effect on accuracy may be due to the failure to choose tests which are complex enough to measure exercise-induced changes in accuracy of performance.” abstract, p. 338. In the present study we assess the impact of acute physical fatigue in professional rugby players with a task that requires both speed of processing and accuracy for adequate performance. We chose to study professional rugby players because this sport has a very high physical demand yet the players are continuously required to process complex dynamic scenes for making appropriate decisions on the field.

## Methods

We tested 22 professional rugby players (age M = 20.54, SD = 1.5; height M = 184. cm, SD = 6.89; weight M = 100.3 Kg, SD = 13.0) from the Top 14 French Professional Rugby League separated into a trained and an untrained group. The trained group underwent 15 training sessions using the CORE program of the NeuroTracker^™^ system (neurotracker.net) distributed by CogniSens Inc. and the untrained group had no prior training. Testing for all conditions was done sitting down on a stationary exercise bicycle and the processing speed threshold scores for the trained group were similar to what was previously obtained when sitting in a regular chair for this population (3). The untrained participants where tested for 3 CORE sessions prior to the fatigue manipulation. The first CORE sequence was used to familiarize the athletes to this task and the baseline value used was the average of the last two CORE sessions. The baseline for the trained group was also the last two values of their 15 CORE sessions (session 14 & 15). For both groups, once the players achieved 80% of the maximum heart rate (80% HRmax) capacity (see methods) they were asked to pedal at a fixed speed while pedal resistance was manipulated to maintained their heart rate at 80% HRmax while the heart rate was monitored. At the same time they executed a CORE session on the NeuroTracker system.

## NeuroTracker task

The task has been described in previous papers (3, 4). The trained group underwent 15 sessions separated over a minimum of five different days using the NeuroTracker^™^ CORE program distributed by CogniSens Athletics Inc., a technology that has been licenced from the Université de Montréal. Each session lasted around 8 minutes and the subjects were not allowed to train for more then three sessions in a given day. The basic trial sequence is presented in Figure 1 and comprises of 5 steps (see legend). In the first step, 8 spheres are randomly positioned in the virtual cube and remain stationary. The next step is the indexation phase where 4 of the 8 spheres are randomly selected and index by changing color and surrounded by a white hallo for a second then they return to their original color and shape undistinguished from the others. The third phase is the movement phase where the spheres move in a straight path but in random directions, bouncing off and redirected when in contact with other spheres or the virtual cube walls (see www.neutracker.net for a demonstration). After 8 seconds of movement the spheres halt. Each sphere then contains a number and the observer has to recall the spheres that were originally indexed in phase 2 by calling out the numbers corresponding to the targets. The size of the virtual cube containing the spheres is 46 degrees of visual angle at the level of the screen. The untrained group did three thresholds on a separate day from testing. All testing prior to the fatigue manipulation was done sitting on the exercise stationary bicycle positioned in the testing room in front of the screen. The geometrical mean of the last two session scores was used as the comparative baseline measure. This was done to allow the untrained group at full 8-minute training session to familiarize them with the procedure. After a single trial (Figure 1), if the subject got all 4 indexed spheres correct the speed went up for the next trial. If at least one sphere was missed the speed slowed down on the next trial (1 up 1 down staircase) so on and so forth until a threshold was achieved (3). All subjects gave the answers verbally and an experimenter recorded the answers on a keyboard. This study was approved by the ethics board of the Université de Montréal.

**Figure 1:**
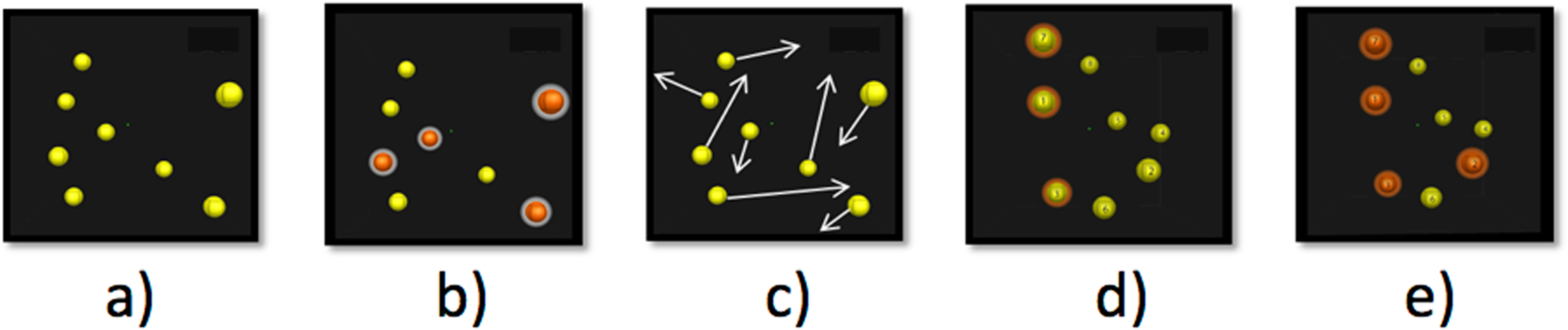
Five steps of the NeuroTracker CORE task^1^ a) <u>presentation phase</u> where 8 spheres are shown in a 3D volume space, b) <u>indexing phase</u> where 4 spheres (targets) change colour (red) and are highlighted (hallo) for 1 second, c) <u>movement phase</u> where the targets indexed in stage b return to their original form and colour and all spheres move for 8 seconds crisscrossing and bouncing off of each other and the virtual 3D volume cube walls that are not otherwise visible, d) <u>identification phase</u> where the spheres come to a halt and the observer has to identify the 4 spheres originally indexed in phase b. The spheres are individually tagged with a number so the observer can give the number corresponding to the original targets, and e) <u>feedback phase</u> where the subject is given information on the correct targets.

## Exercise

On the exercise manipulation day, the players warmed up on the exercise stationary bicycle (schwinn evolution-sr) for about 2 minutes while the trainer continuously monitored the heart rate with a Polar heart monitor (Kalenji, cw400 coded). They were specifically instructed to pedal at 90 cycles per minute and guided with the use of a metronome (iPhone app). The pedal resistance was adjusted to obtain 80% HRmax. Once this level was attained, the NeuroTracker test started while the subject maintained the pedaling rate and the 80% HRmax was kept constant by the experimenter by continuous monitoring and adjustment of pedal resistance.

## Results

Figure 2 shows the baseline data prior to the physical exercise versus the CORE speed threshold obtained while at 80% HRmax. A 2×2 between (group) within (fatigue) ANOVA was performed showing a significant group by fatigue interaction (F(1,20) = 4.47; p = 0.047) along with significant main effects of fatigue (F(1,20) = 8,41; p = 0.009) and group (F(1,20) = 35.44; p < 0.001). The data clearly show that maintaining an 80% HRmax level had a significant negative impact on the speed thresholds for processing complex dynamic scenes in the untrained group as compared to the trained group. The speed decrease was present in the untrained group even though the repeated testing session should have increased their values significantly from the baseline as the 4^th^ session speed scores are usually undergoing steep learning functions in this population of professional athletes with an average of 0.20 units increase (3). On the other hand the 16^th^ training session used for our trained group normally would show shallow learning functions (3) with less then 0.02 units increase, placing the anticipated training level well within the standard error range presented here. An asterisk is shown in Figure 2 to illustrate how large the difference is between what would normally be obtained from a training increase for the same session versus what we observe in the 80% HRmax fatigue condition. The drop of performance at 80% HRmax is of 0.38 units. Adding the 0.20 unit increase that would normally be obtained for the same training session with the 0.38 drop at 80% HRmax results in a differential of 0.58 speed units between what is obtained and what would be expected if the physical exercise had no impact on the untrained group for processing this task. In percentage decrease, the change in performance for the trained group is only of 0.03% at 80% HRmax while the change is 21% for the untrained group. Furthermore, if one accounts for the decrement caused by fatigue with the anticipated scores of athletes from the same population as mentioned above (0.02 increase at training session 16 and 0.20 at training session 4), the difference remains at 0.03% for the trained group while the untrained group shows a 30% decrement in performance.

**Figure 2:**
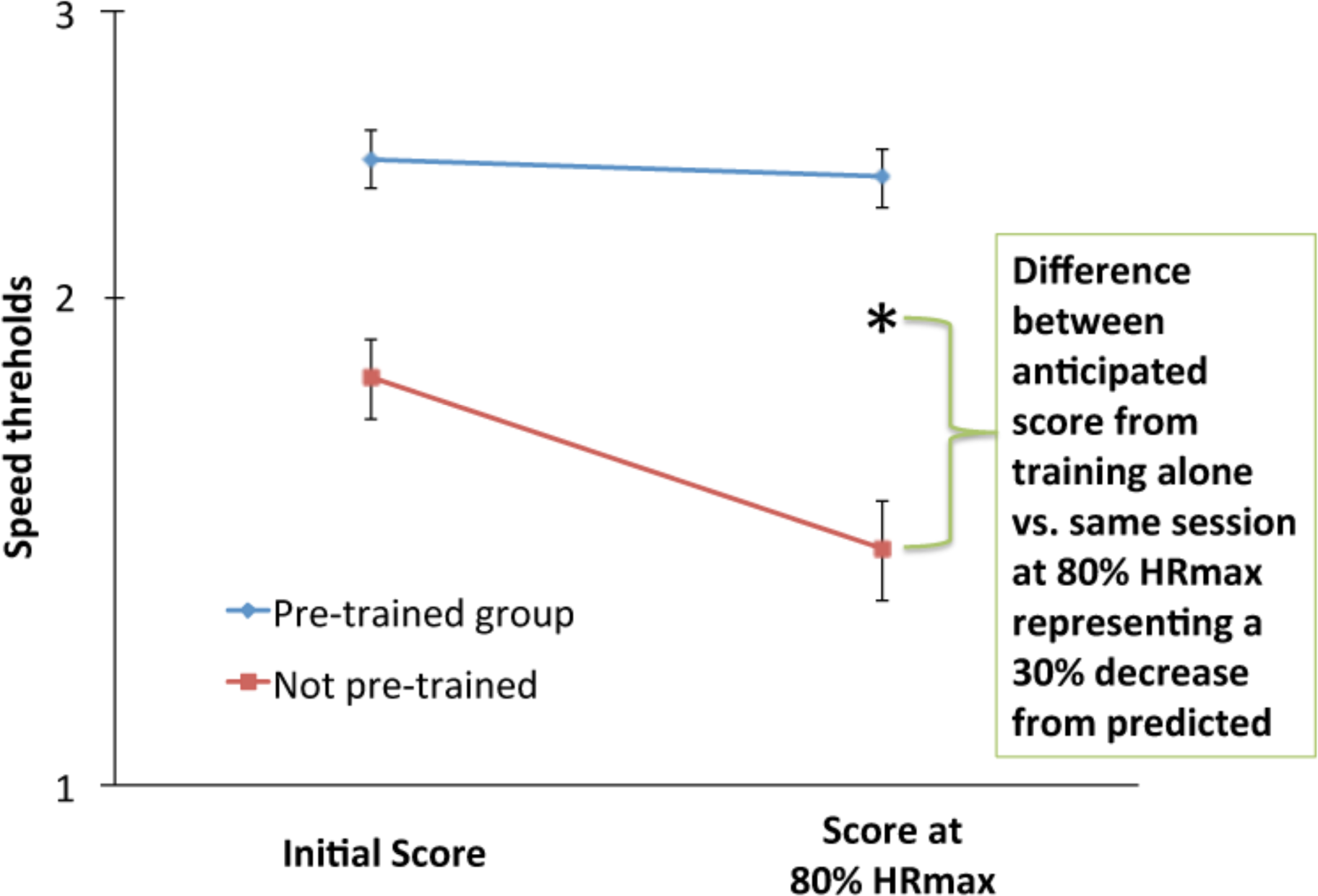
Geometrical NeuroTracker CORE speed threshold means for 22 individuals on a log scale for previously trained and untrained rugby professional players as a function of baseline versus 80% HRmax. Error bars represent SEM.

## Discussion

The results show that acute physical exercise can negatively impact the capacity for processing complex dynamic visual scenes and that this effect can be dramatically reduced if the professional player has had prior perceptual-cognitive training. Our results are quite novel on different levels. We show that this complex dynamic scene task is demanding enough to measure a significant negative impact of isolated acute physical fatigue on performance in professional athletes, which in its own right is novel. We also show that prior training will mitigate the negative impact of acute physical fatigue on the ability for processing complex dynamic scenes. This is critical for professional players who are always striving to “read the game” efficiently and therefore make appropriate decisions on the field even when under acute physical activity. It is therefore conceivable that optimal sports training regimes should incorporate perceptual-cognitive training protocols of the likes that are presented here. It has been shown that perceptual-cognitive training for the NeuroTracker task transfers into benefits for previously untrained tasks of social relevance such as biological motion perception (5) critical for high level sports performance (1, 10). It was also demonstrated that measures on this task are predictive of sports decision making metrics in professional athletes (6) and that training on this task generates transfer in the field for sports related decisions during play (7). It is important to note that we have only isolated the physical fatigue component in the present study, as minimal mental resources are required for professional athletes to maintain heart rate at this fixed level. It would be very interesting to determine the extent of what is shown here when additional sports specific precision motor control requirements are necessary. It was previously shown that motor requirements may have a significant impact on the learning function of the dynamic scene task presented here (4) yet we still lack information on the detrimental effects of combined fatigue, precision motor control tasks and mental decision making on the field. Future studies on the potential benefit of training regimens on these abilities would be very interesting. Another avenue of interest would be to measure the recovery of these athletes at different time scales after the exercise. Unfortunately our time with the professional athletes was limited and did not allow us to repeat the NeuroTracker measure during recovery. It would be interesting to determine whether the NeuroTracker scores would come back to their initial levels prior to the fatigue task or even be improved due to an increased arousal state (2).

In conclusion, we show that acute physical fatigue has a detrimental impact on perceptual-cognitive processing as measured by the 3D-MOT task in professional athletes and that prior perceptual-cognitive training can dramatically reduce this negative consequence hence, reduce decision-making errors so critical for elite athletic performance.

## Acknowledgements

This work was supported by a Natural Sciences and Engineering Research Council of Canada discovery grant. We would like to thank Dr. Leonard Zaichkowsky for helpful discussions and Isabelle Legault for help with the analysis.

## Author Contribution Statement

JF wrote the manuscript text. SB executed the experimental protocols. SB and JF designed the experiments did the analysis and prepared the figures.

## Additional information

(Competitive financial interests) JF is director of the Visual Faubert Laboratory at the University of Montreal and he is the Chief Science Officer of CogniSens Athletics Inc. who produces the commercial version of the NeuroTracker used in this study. In this capacity, he holds shares in the company. During this study, SB was a PhD student in Visual Neurosciences at the School of Optometry of Université de Montréal.

